# A new *Spirodela polyrhiza* genome and proteome reveal a conserved chromosomal structure with high abundances of proteins favoring energy production

**DOI:** 10.1101/2020.01.23.909457

**Authors:** Alex Harkess, Fionn McLoughlin, Natasha Bilkey, Kiona Elliott, Ryan Emenecker, Erin Mattoon, Kari Miller, Kirk Czymmek, Richard Vierstra, Blake C. Meyers, Todd P. Michael

## Abstract

Duckweeds are a monophyletic group of rapidly reproducing aquatic monocots in the Lemnaceae family. *Spirodela polyrhiza*, the Greater Duckweed, has the largest body plan yet the smallest genome size in the family (1C = 150 Mb). Given their clonal, exponentially fast reproduction, a key question is whether genome structure is conserved across the species in the absence of meiotic recombination. We generated a highly contiguous, chromosome-scale assembly of *Spirodela polyrhiza* line Sp7498 using Oxford Nanopore plus Hi-C scaffolding (Sp7498_HiC) which is highly syntenic with a related line (Sp9509). Both the Sp7498_HiC and Sp9509 genome assemblies reveal large chromosomal misorientations in a recent PacBio assembly of Sp7498, highlighting the necessity of orthogonal long-range scaffolding techniques like Hi-C and BioNano optical mapping. Shotgun proteomics of Sp7498 verified the expression of ∼2,250 proteins and revealed a high abundance of proteins involved in photosynthesis and carbohydrate metabolism among other functions. In addition, a strong increase in chloroplast proteins was observed that correlated to chloroplast density. This Sp7498_HiC genome was generated cheaply and quickly with a single Oxford Nanopore MinION flow cell and one Hi-C library in a classroom setting. Combining these data with a mass spectrometry-generated proteome illustrates the utility of duckweed as a model for genomics- and proteomics-based education.

## Introduction

Duckweeds are the fastest reproducing flowering plants, with some species capable of dividing in as little as 18 hours, under optimal conditions (Ziegler *et al*., 2015). Found on every continent except Antarctica, the Lemnaceae family of duckweeds is taxonomically divided into 37 species across five genera: *Spirodela, Lemna, Landoltia, Wolffia*, and *Wolffiella*. They are strictly aquatic, typically floating on slow-moving freshwater.

*Spirodela polyrhiza* has the smallest known duckweed genome size at 158 Mb (Wang *et* al., 2011). The initial *Spirodela* reference generated by the Department of Energy Joint Genome Initiative (DOE-JGI) was based on clone 7498 (Sp7498) collected from Durham, North Carolina USA (35N 75W) that was received directly from Elias Landolt, an early pioneer of duckweed taxonomy, physiology, and anatomy (Urbanska *et al*., 2013). The initial Sp7498 genome assembly was based on 454 pyrosequencing reads and BAC-end sequencing. The resulting draft assembly was sufficiently contiguous to reveal that while *Spirodela* retains a core set of plant genes, it shows a reduction in most gene families resulting in a total of 19,623 protein coding genes (Wang *et al*., 2014). At the time this was the fewest number of genes found in a flowering plant genome, perhaps consistent with its greatly reduced body plan and rapid growth. Afterwards, the sea-grass genome *Zostera marina* was sequenced and also had a reduced protein coding gene set at 20,450, which suggests that a reduced gene set may be a feature of aquatic plants (Olsen *et al*., 2016). However, the Sp7498 genome assembly only covered 90% of the expected genome size, with 10.7% of the assembly in gaps filled with Ns. Instead of being resolved into the expected 20 chromosomes, it was anchored onto 32 pseudo-molecules (Wang *et al*., 2014).

*Spirodela* species form specialized, dormant winter buds called turions that accumulate high levels of starch (40 to 70% dry weight) and sink to the bottom of the water source where they rest, surviving on starch reserves, until favorable growing conditions stimulate the resumption of vegetative growth. A second clone 9509 (Sp9509) from Lotschen, Stadtroda Germany (50N 11W) was sequenced and assembled to help understand turion formation (Michael *et al*., 2017). Sp9509 was sequenced using a combination of Illumina short read libraries and scaffolded into 20 chromosomes using BioNano Genomics Optical maps. In combination with extensive expression (RNA-seq) and DNA methylation (bisulfite-seq) analysis, the minimal set of genes was confirmed, which also revealed that *Spirodela* has the lowest levels of genome-wide DNA methylation of any flowering plant examined (Michael *et al*., 2017). The Sp9509 assembly was later updated using Oxford Nanopore single molecule long read sequencing and the chromosomal structure was validated using orthogonal techniques, which resulted in the most accurate and contiguous *Spirodela* genome to date (Hoang *et al*., 2018).

Recently, the Sp7498 genome was updated using Pacific Biosciences (PacBio) Sequel reads (An *et al*., 2019). The resulting assembly is an order of magnitude less contiguous than Sp9509 with a contig N50 length of 0.83 Mb compared to Sp9509 at 2.87 Mb (Table 1), and does not leverage information from chromosome-scale technologies (Michael *et al*., 2017; Hoang *et al*., 2018), such as BioNano optical mapping or Hi-C chromatin conformation sequencing. To identify any possible mis-assemblies in the latest PacBio Sp7498 genome assembly, and more broadly to assess genome stability across the *Spirodela polyrhiza* species, we generated a single MinION flow cell of Oxford Nanopore long-read data for this line, as well as Phase Genomics Hi-C data, as part of a graduate seminar course at Washington University in St. Louis. As a novel strategy to more accurately map exomes, we also subjected Sp7498 to shotgun proteome analysis via tandem mass spectrometry (MS/MS) following liquid chromatographic separation of trypsinized soluble proteomes, which helped verify the expression of ∼2,250 proteins. We combined these data to present a high-quality genome assembly and proteome for Sp7498, and we highlight the usefulness of duckweeds in a classroom setting on genomics and proteomics.

**Table 1:** Assembly statistics for 7498 and 9509 genomes. The genome assemblies were compared using quast to generate basic statistics. PacBio, Pacific Bioscience; Oxford, Oxford Nanopore; HiC, high throughput chromatin conformation capture; BioNano, BioNano Genomics optical mapping; mcFISH, multi-color FISH; N50 length, the length of the contig at half of the assembly; L50, the number of contigs at half the length of the assembly; N’s, gap in the assembly filled with N to represent unknown bases.

|  | Sp7498_PNAS | Sp7498_PNAS | Sp7498_HiC | Sp7498_HiC | Sp9509 | Sp9509 |
| --- | --- | --- | --- | --- | --- | --- |
| sequencer | PacBio | PacBio | Oxford | Oxford | Oxford | Oxford |
| Chromosome | BAC | BAC | HiC | HiC | BioNano/mcFISH | BioNano/mcFISH |
|  | Scaffolds | contigs | Scaffolds | contigs | Scaffolds | contigs |
| Total length (bp) | 138,536,245 | 138,525,388 | 138,493,532 | 138,486,032 | 138,592,155 | 138,570,896 |
| Largest contig (bp) | 10,795,902 | 3,236,013 | 11,339,703 | 8,229,153 | 11,560,055 | 7,326,273 |
| contigs (#) | 134 | 384 | 24 | 99 | 20 | 95 |
| N50 length (bp) | 7,645,982 | 837,937 | 7,689,391 | 3,339,466 | 7,949,387 | 2,868,147 |
| L50 (#) | 8 | 52 | 8 | 15 | 8 | 16 |
| N's (#) | 10,857 | 0 | 7,500 | 0 | 21,268 | 9 |

## Methods

### Sp7498 Oxford Nanopore genome sequencing, assembly, and annotation

*Spirodela polyrhiza* line Sp7498 tissue was collected from the Rutgers Duckweed Stock Cooperative (RDSC; http://www.ruduckweed.org/) and grown in Hoaglands No. 2 Basal Salt Mixture (1.6 g/L) at 25C under 16 hour days. High molecular weight DNA was isolated from 1.5 g of flash-frozen whole-plant *Spirodela polyrhiza* 7498 tissue using 10 mL CTAB (100mM Tris-HCl pH 8.0; 1.4 M NaCl; 20 mM EDTA; 2% cetyl trimethyl ammonium bromide (CTAB); 2% polyvinylpyrrolidone (PVP); 0.2% 2-mercaptoethanol) in a 65C waterbath for 45 minutes. DNA was purified twice with 1 volume of 24:1 chloroform:isoamyl alcohol, precipitated with isopropanol and spooled out on a glass shepherd’s crook, then treated with 3ul RNase A (20 ug/mL) at 37C for 30 minutes, then left on a benchtop at room temperature overnight. DNA was purified again with 24:1 chloroform:isoamyl alcohol, and resuspended in TE buffer on a benchtop at room temperature for 48 hours. Approximately 1 ug of DNA measured by Qubit dsDNA BR were used as input to the LSK-109 library kit, loaded onto a 9.4.1 flow cell and sequenced for 48 hours. Raw signal data were basecalled using Guppy v3.1.5 with the “flip-flop” algorithm. Reads were assembled with minimap2 (2.17-r941) (Li, 2018) and miniasm (0.2-r128) (Li, 2016) with default options, then error-corrected with three rounds of Racon (Vaser *et al*., 2017) and one round of Medaka (https://github.com/nanoporetech/medaka).

Hi-C data were generated from frozen whole plant line 7498 tissue sent to Phase Genomics (Seattle, WA) using the Sau3IA cut-site and assembled into pseudomolecules with Proximo software. Gene annotations were lifted over from Sp9509 to the Sp7498_HiC assembly. The resulting assembly and annotation was syntenically compared to Sp9509 and Sp7498_PNAS using CoGE SynMap with default DAGChainer options (-D 20 -A 5) (Lyons, 2008). For more detailed statistics on misassemblies, mismatches, and indels, QUAST v5.0.2 with default options was used (Gurevich *et al*., 2013). LTR retrotransposons were annotated in all three genomes using default parameters in GenomeTools LTRharvest (v1.5.8) (Ellinghaus *et* al., 2008).

### Mass Spectrometry proteome profiling

Proteins were extracted at 4°C from pulverized *Spirodela polyrhiza* 7498 and *Arabidopsis thaliana* Col-0 leaf tissue into 50 mM HEPES (pH 7.5), 5 mM Na_2_EDTA, 2 mM dithiotreitol, and 1X plant protease inhibitor cocktail (Sigma-Aldrich). The samples were further homogenized using a Pyrex Potter-Elvehjem tissue grinder (Fisher Scientific) and clarified by centrifugation at 16,000 x *g*. 150 µl of the protein extract was precipitated in 4:1:3 (v/v) methanol/chloroform/water and collected by centrifugation. The resulting pellet lyophilized to dryness, and resuspended into 100 µl of 8 M urea, reduced for 1-hr at 22 °C with 10 mM dithiothreitol, followed by alkylation with 20 mM iodoacetamide for 1-hr. The reaction was quenched with 20 mM dithiotreitol and diluted with with 900 µl of 25 mM ammonium bicarbonate to reduce the urea concentration below 1.5 M, and digested overnight at 37°C with 0.5 µg of sequencing-grade modified porcine trypsin (Promega). The resulting peptides were lyophilized to a final volume of ∼250 µl, acidified with 0.5% v/v trifluoroacetic acid (pH<3.0), and desalted and concentrated using a 100 µl Bond Elut OMIX C18 pipette tip (Agilent Technologies) according to the manufacturer’s instructions. The peptides were eluted in 50 µl of 75% acetonitrile and 0.1% acetic acid, lyophilized, and resuspended in 20 µl 5% acetonitrile and 0.1% formic acid.

Nano-scale LC separation of the tryptic peptides was performed using a Dionex Ultimate 3000 Rapid Separation system equipped with a 75 µm x 25 cm Acclaim PepMap RSLC C18 column (Thermo Fisher Scientific), in combination with a 2-hr linear 4% to 36% acetonitrile gradient in 0.1% formic acid and a flow rate of 250 nl/min. Eluted peptides were analyzed online by a Q Exactive Plus spectrometer (Thermo Fisher Scientific) in the positive ESI mode. Data-dependent acquisition of full MS scans (mass range of 380-1500 m/z) at a resolution of 70,000 was collected, with the automatic gain control (AGC) target set to 3 × 10^6^, and the maximum fill time set to 200 msec. High-energy collision-induced dissociation fragmentation of the 15 strongest peaks was performed with an intensity threshold of 4 × 10^4^ counts and an isolation window of 3.0 m/z, and excluded precursors that had unassigned, +1, +7, +8, or >+8 charge states. MS/MS scans were conducted at a resolution of 17,500, with an AGC target of 2 × 10^5^ and a maximum fill time of 100 msec. Dynamic exclusion was performed with a repeat count of 2 and an exclusion duration of 30 sec, while the minimum MS ion count for triggering MS/MS was set to 4 × 10^3^ counts. Each sample was analyzed in quadruplicate to enable broad coverage; the first two runs were performed without an exclusion list, while the third and fourth runs were performed with an exclusion list containing the 5,000 most abundant peptides that were detected in the first two runs, to increase sample coverage and maximize suppression of abundant peptides. Raw MS2 files from all four runs were merged, resulting in two technical replicates per sample (McLoughlin *et al*., 2018).

The resulting MS/MS datasets were queried by Proteome Discoverer (version 2.0.0.802; Thermo Fisher Scientific) against the *S. polyrhiza* Sp7498 (Spolyrhiza_290_v2.protein.primarytranscriptonly.header.fa; http://phytozome.jgi.doe.gov) and *Arabidopsis* (TAIR10_PEP_20101214_UPDATED.fa; http://www.arabidopsis.org) predicted proteome databases and a list of common protein contaminants. Peptides were assigned by SEQUEST HT (Eng *et al*., 1994), allowing a maximum of 1 missed tryptic cleavage, a minimum peptide length of 6, a precursor mass tolerance of 10 ppm, and fragment mass tolerances of 0.02 Da. Carbamidomethylation of cysteines and oxidation of methionine were specified as static and dynamic modifications, respectively.

The target false discovery rate (FDR) of ≤0.01 (strict, high confidence) and 0.05 (relaxed, medium confidence) were used as validation for peptide spectral matches (PSMs), protein modifications are only reported at high PSM confidence (FDR 0.01) and peptides and protein grouping was performed excluding all protein groups that are not strictly necessary to explain the identified peptides (strict). Label-free quantification was obtained in Proteome Discoverer™ as previously described (Silva *et al*., 2006) with a minimum Quan value threshold set to 0.0001 using unique peptides, and “3 Top N” peptides used for area calculation (Silva *et al*., 2006). Three biological replicates were each analyzed in quadruplicate, and the resulting values were averaged, which resulted in 2,163 master protein identifications. Data was log_2_ transformed and missing values were imputed while assuming a normal distribution using a width distribution shrinkage of 0.3 and a downshift of 1.8 standard deviations if necessary using the Perseus computational platform (Tyanova *et al*., 2016). Peptides were analyzed and significances were determined by plotting the datapoints in a volcano plot using a t-test with a number of randomizations of 250 with a FDR of 0.05 and an S0 of 0.1. GO enrichments were identified by the AgriGO analysis toolkit (Tian *et al*., 2017). GO-annotation categories shown here were selected based on their uniqueness, *p*-value significance, and degree of completeness.

### Confocal microscopy and chloroplast density

We used mature, fully-expanded leaves of *Arabidopsis thaliana* Col-0 and *S. polyrhiza* Sp7498 mother frond to image and quantify chloroplast density and distribution. Confocal microscopy was performed by syringe infiltration of *A. thaliana* leaf and *S. polyrhiza* fronds with Perfluoroperhydrophenanthrene (Sigma-Aldrich, Cat # 58919) to fill air spaces and allow deep imaging into the mesophyll by confocal microscopy (Littlejohn et al., 2014). 3D volumes were captured on a Leica SP8-X using HC PlanApochromat 63X Water Immersion Objective Lens with the pinhole set to 1 Airy unit and z-step of 0.6 microns. 405 and 649 laser excitation with the spectral prism set to collect 414nm - 516nm and 658 - 768nm emission for imaging cell wall and chloroplast autofluorescence, respectively. 3D volumes were processed in FIJI (ImageJ version 1.51w) by thresholding background in the red channel (representing chloroplast autofluorescence) and averaging the calculated percent area of chloroplasts in each volume with slices extending from the epidermis to the mesophyll within the plant tissue.

## Results

To generate a high quality genome assembly for Sp7498, we sequenced long-read DNA sequence on one Oxford Nanopore MinION 9.4.1 flow cell. After filtering poor quality “failed” reads with the Guppy base caller, we generated 1.4 million clean single molecule nanopore reads with an N50 of 13.5 kb and total length of 6.4 gigabases, covering approximately 46X coverage of the roughly 138 Megabase *S. polyrhiza* 7498 genome. Genome assembly with miniasm yielded a high quality contig assembly with a contig N50 of 3.34 Mb (Figure 1A). The total length of the assembly was 138.49 Mb, which is highly congruent with all existing Sp7498 assemblies regardless of technology (Table 1).

**Figure 1:**
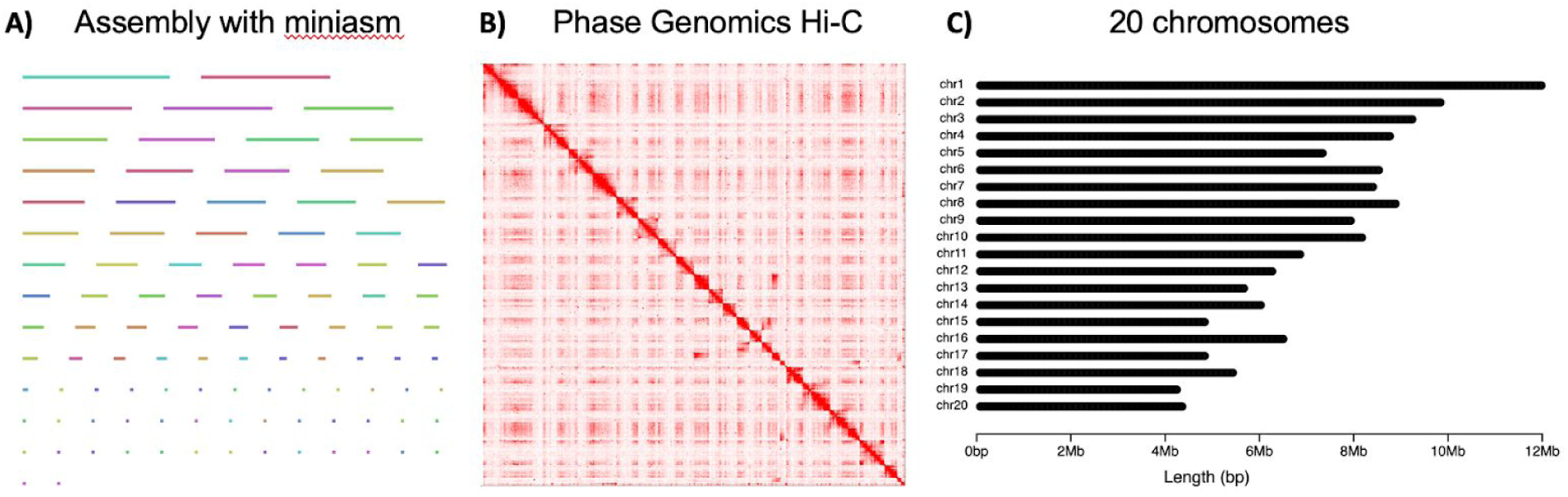
Assembly and scaffolding strategy overview for the Sp7498 genome. A) Contigs were assembled from raw Oxford Nanopore reads using miniasm to generate 99 contigs. Contigs were polished with three iterations of Racon, followed by one iteration of Medaka, and visualized with Bandage. B) Chromosome scaffolding using HiC links. A Phase Genomics Hi-C library was sequenced, followed by Proximo scaffolding and manual polishing in JuiceBox into chromosome pseudomolecules. C) Chromosome naming and length distribution. All 20 chromosomes were named according to synteny against the Sp9509 assembly.

To scaffold the contig assembly into chromosomes, a Phase Genomics Hi-C library was generated and sequenced to 115,741,530 read pairs. Further scaffolding and polishing with the Phase Genomics Hi-C links using Proximo identified no misjoins in the assembly, and combined the assembly into 20 chromosomes plus four unplaced scaffolds (Figure 1B, Figure 1C). The four unplaced scaffolds total 2.8 Mb, or 2.0% of the total assembly length. The canonical telomere repeat (5’-TTTAGGG-3’) is the same in Sp7498_HiC as identified in Sp9509 (Michael *et al*., 2017). All 20 chromosomes in Sp7498_HiC contain a telomere repeat at the distal tip of at least one arm of the chromosome, and 12 chromosomes have telomere repeats at the distal tips of both arms (Supplemental Figure 2).

**Figure 2:**
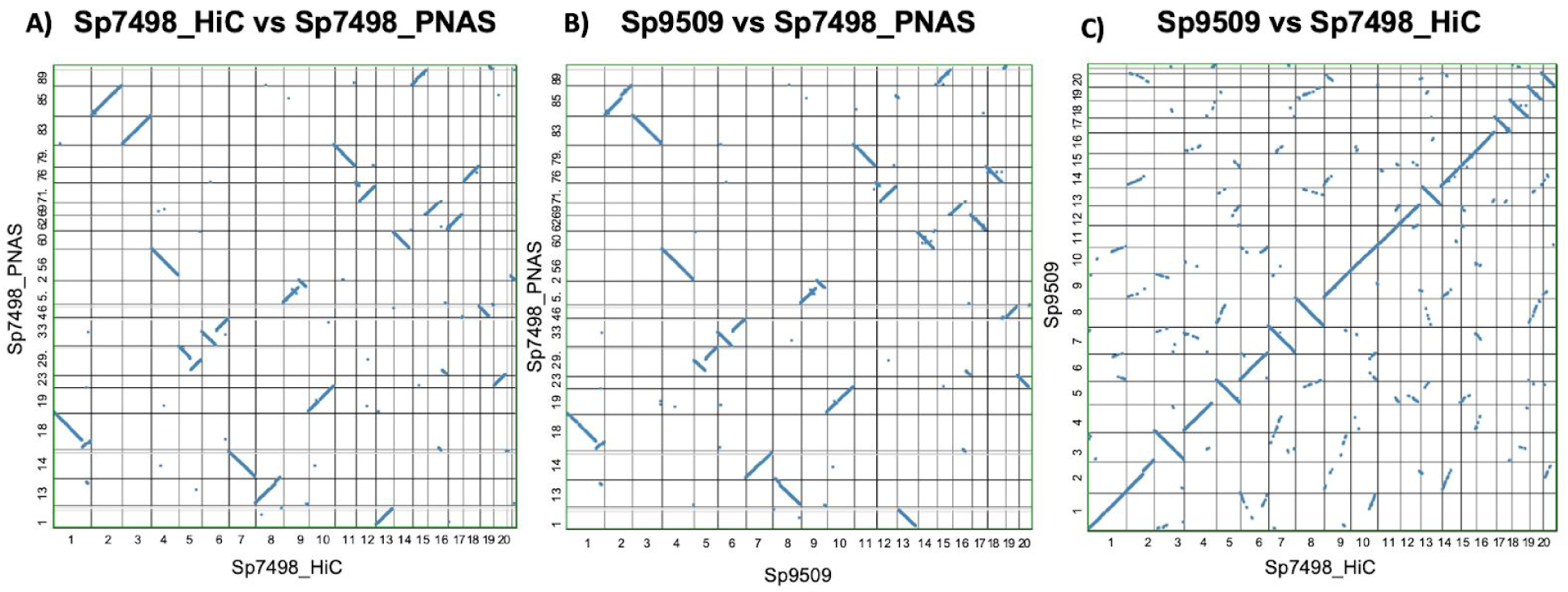
Syntenic dotplot comparisons of Sp7498 and Sp9509 genomes. All comparisons were performed in CoGE using default SynMap parameters. Any comparison involving Sp7498_PNAS was performed using gene predictions lifted over from Sp9509 since no gene annotations are publicly available. A) Sp7498_HiC vs Sp7498_PNAS, B) Sp9509 vs Sp7498_PNAS, and C) Sp9509 vs Sp7498_HiC.

We then tested the degree of synteny of our Sp7498_HiC assembly against the recently published Sp7498_PNAS genome, which should be genotypically identical. The assembled genome sizes are nearly the same, differing by only 42,713 nt more in the Sp7498_HiC assembly. Across the whole genome, GC content is nearly identical as well (Supplemental Figure 1) A synteny dot-plot reveals several major discrepancies between the two assemblies, including several whole chromosome arm inversions (Figure 2A). Across the aligned length of both genomes, 689 misassemblies (breakpoints) were identified when using Sp7498_HiC as the reference. These include 8 inversions, 96,774 single nucleotide mismatches and 315,762 indels, 96% of which are less than 6 nt in size. Nearly 1.6 Mb of sequence in the Sp7498_HiC assembly do not share an alignment with the Sp7498_PNAS genome.

Next, we tested the level of conservation across two unique clones of *Spirodela polyrhiza*, Sp7498 and Sp9509. Both of these genomes (Sp7498_HiC and Sp9509) were assembled with similar methods, based on Oxford Nanopore MinION 9.4.1 flow cells and assembly with miniasm. Overall the two genomes are highly syntenic across the whole genome, and share nearly all of the same contig order and orientation for all 20 chromosomes (Figure 2C). Both the Sp7498_HiC and Sp9509 genomes share the same large structural variation disagreements with the Sp7498_HiC genome (Figure 2B). Of the 22,605 peptides annotated in the Sp9509 genome, we were able to lift over 20,530 peptides onto the Sp7498_HiC genome.

We next asked whether repeat content variation could explain some of this variation between genome assemblies. In all three assemblies (Sp7498_HiC, Sp7498_PNAS, Sp9509), repeat content of LTR retrotransposons is low, with between 1,511 and 1,647 annotations (Figure 3). Sp7498_PNAS contains the greatest number of LTR annotations (N=1,647), whereas Sp9509 has the fewest (N=1,511). We compared these three long-read assemblies to the first *Spirodela polyrhiza* genome (Sp7498_v3), which was produced with a combination of 454 pyrosequencing, Sanger reads, and BAC-end sequencing. The Sp7498_v3 assembly shows a marked decrease in total number of LTR annotations, as well as a reduction in young (ie. high percent identity of LTR ends) retrotransposons in the assembly.

**Figure 3:**
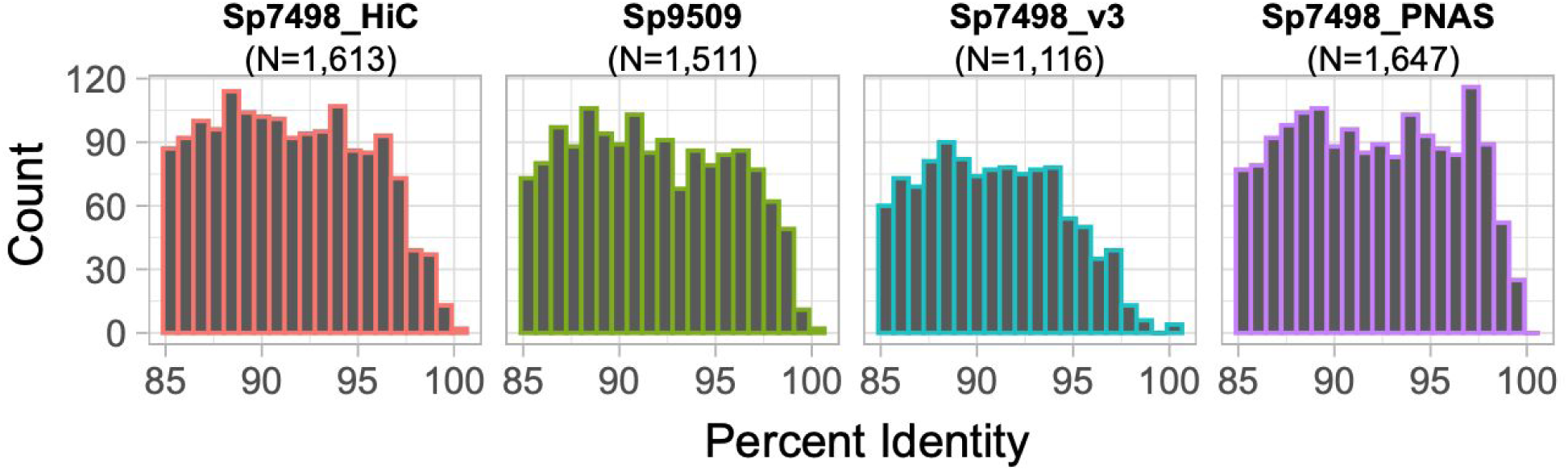
Comparison of LTR retrotransposon content and estimated insertion timing across four *Spirodela polyrhiza* genome assemblies. Percent identity was calculated by pairwise comparisons of LTR ends for each full-length LTR retrotransposon.

To validate the expression of a plethora of protein coding regions we conducted a protein mass spectrometric analysis on the total protein extracts of *Spirodela* which resulted in the detection of 2,289 proteins. Although the protein coding regions were generally uniformly mapped to all chromosomes, clusters were observed on chromosomes 4, 6, 7, 10 and 13 (Figure 4A). In contrast, chromosomes 15 and 17 showed a reduced amount of confirmed protein coding regions, indicating that these chromosomes do not contain many abundantly expressed proteins.

**Figure 4:**
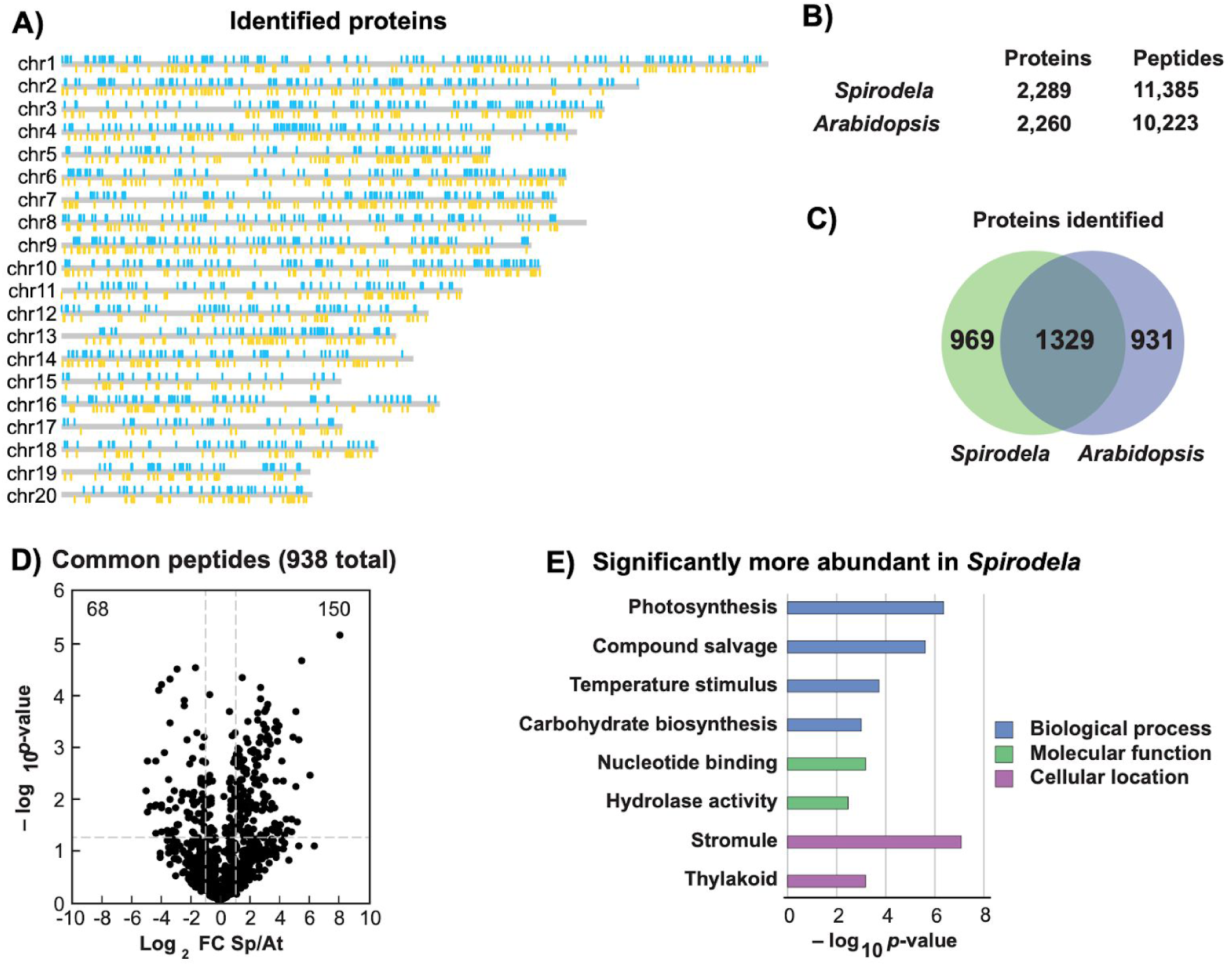
Analysis of the *Spirodela polyrhiza* proteome. Proteins were identified and quantified from a total soluble extract. A) Confirmed protein coding regions are highlighted on the genome, top strand in blue and bottom strand in yellow. B) Crude protein extract of *Spirodela* and *Arabidopsis* resulted in the identification of ∼2250 proteins. C) Venn diagram displaying the overlap of detected protein homologues in *Spirodela* and *Arabidopsis*. D) Quantitative analysis of conserved peptides between *Spirodela* and *Arabidopsis*. 938 conserved peptides were identified and quantified using MS1 precursor intensity. Significance in abundance was called at p<0.05, FC>2x (n=3). E) Proteins that harbour two significantly affected peptides were selected and gene ontology analysis was conducted. P-values for significant categories were -log transformed and plotted.

To pinpoint characteristics of the *Spirodela* proteome, a comparative analysis was conducted with the model species *Arabidopsis thaliana*. Mass spectrometric analysis resulted in the identification of more than 11,000 and 10,000 unique peptides for *Spirodela* and *Arabidopsis*, respectively, which were assigned to ∼2,250 protein groups in both species (Figure 4B, Supplemental Tables 1 and 2). Protein homologues were assigned and compared to determine the overlap between the detected *Spirodela* and *Arabidopsis* proteome which resulted in the identification of ∼1,300 commonly detected protein homologues where ∼950 were only detected uniquely for each species (Figure 4C, Supplemental Table 3). Since it is hard to determine if the uniquely detected proteins are biologically relevant or not detected or assigned due to variations in protein coding sequences or differences in protein grouping, we restricted our analysis to perfectly conserved peptides between *Spirodela* and *Arabidopsis*.

To hone in on processes or cellular components that are either over- or underrepresented in *Spirodela*, we selected 935 peptides that were perfectly conserved between *Arabidopsis* and *Spirodela*. MS1 precursor abundance was used to quantify the peptides and significantly abundant peptides were calculated and plotted in a volcano plot (Figure 4D) where 68 peptides were significantly more abundant in *Arabidopsis* and 150 in *Spirodela*. We then selected proteins of which at least two peptides were significantly affected (Supplemental Table 4) and Gene Ontology (GO) analysis was conducted using all the commonly detected proteins as a reference (Figure 4E). Although there were no significantly affected categories in *Arabidopsis*, an overrepresentation of proteins involved in various processes directly related to energy production (*e*.*g*. photosynthesis and carbohydrate metabolism) was observed. The overrepresentation of the energy production related protein clusters is consistent and could contribute to the rapid reproduction rate of *Spirodela*.

To validate the increase of several representative energy production related proteins, we plotted the relative abundance of conserved peptides of *Arabidopsis* and *Spirodela* (Fig. 5A). Among the abundantly expressed proteins in *Spirodela* were Ribulose-bisphosphate carboxylase; RuBisCO (carbon fixation), glyceraldehyde-3-phosphate dehydrogenase; GAPDH A and B (glycolysis), photosystem I reaction center subunit; PSAN (light driven oxidoreductase) and phosphoribulokinase; PRK (Calvin cycle). The increase in associated peptide abundances were consistent for all the selected proteins and indicates a high abundance of various energy related processes in *Spirodela*. Since these processes all exclusively occur in the chloroplast, the increase in the proteins could reflect a high density or enlarged chloroplasts in *Spirodela*. This would be consistent with the increase in chlorophyll content apparent by the darker green color in the protein extracts (Supplemental Figure 3). Therefore, chloroplasts were imaged in *Spirodela* and *Arabidopsis* using a Z-stack of confocal images of a fully expanded plant of Sp7498 (Figure 5B). In comparison to *Arabidopsis, Spirodela* displays a higher number of smaller chloroplasts with less spatial organization (Figure 5B), although the total area of chloroplasts in the leaf is not significantly different (Figure 5C). Altogether, these results suggest that the relatively smaller chloroplasts in *Spirodela* contain more protein per volume, potentially resulting in a very effective energy production strategy compared to *Arabidopsis*.

**Figure 5:**
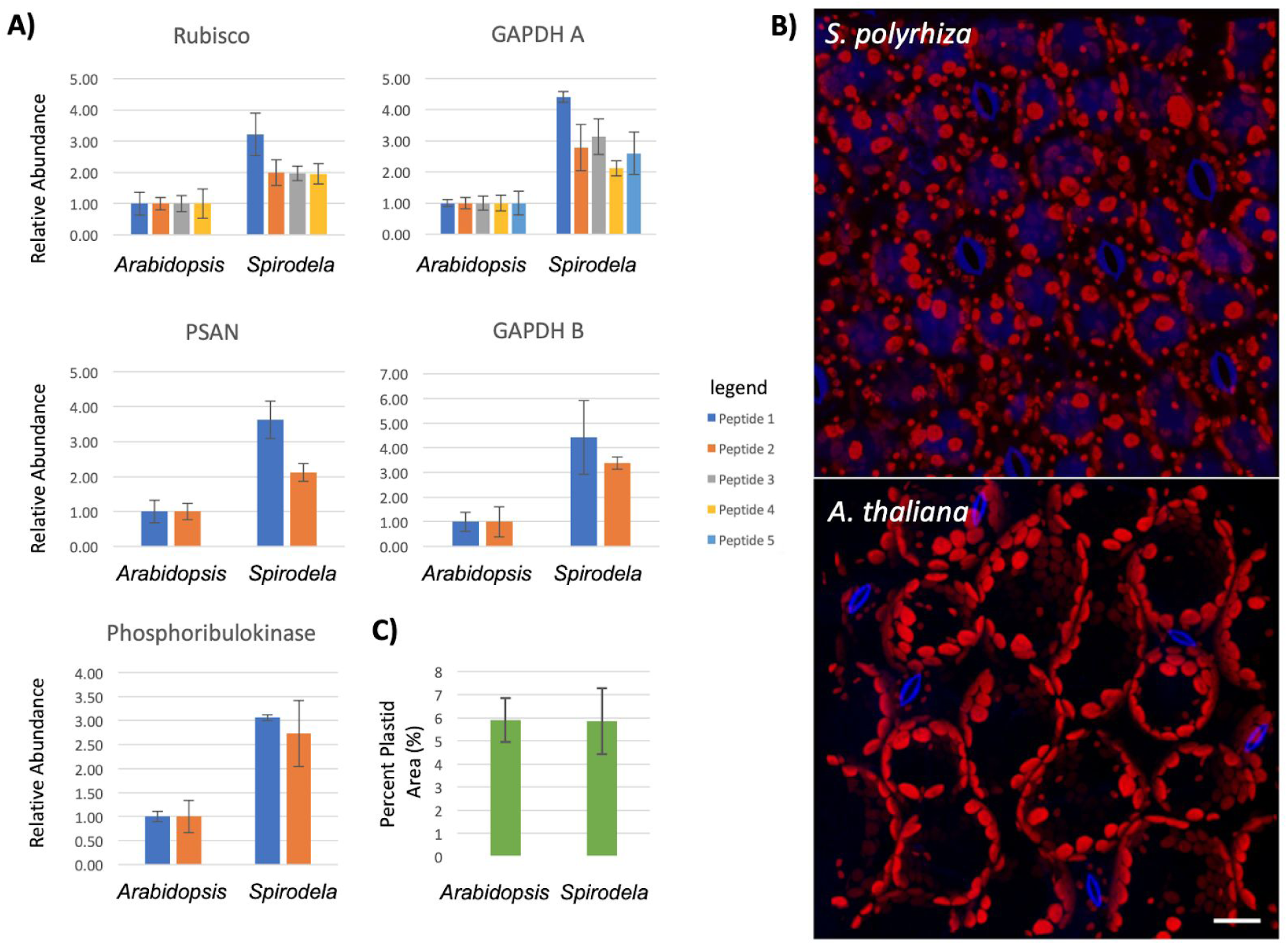
Quantification of peptides from various chloroplast proteins and the chloroplast abundance and distribution. A) Peptide abundances were determined using the MS1-precursor intensity and normalized to the average value obtained from *Arabidopsis*. Each bar represents the average value of a unique and conserved peptide and the error bars represent the standard deviation from the mean. B) Confocal microscopy of *Spirodela* and *Arabidopsis* to measure chloroplast density in 3D stacks. Blue is UV autofluorescence of cell walls and vacuoles, and red is autofluorescence of chlorophyll. Scale bar is 20 μm. C) Measurement of average percent area of chloroplasts through the 3D volumes extending from the epidermis to the mesophyll. The error bars represent the standard deviation from the mean.

## Discussion

Here, we present an independently assembled reference genome using Oxford Nanopore and Phase Genomics Hi-C for *Spirodela polyrhiza* clone 7498 (Sp7498). We draw comparisons to another assembly of the same clone, as well as assess the structural diversity between another clone Sp9509. Our Sp7498 assembly shows large-scale discrepancies compared to a different assembly of the same line, yet high chromosomal conservation compared to Sp9509. The conserved genome structure between Sp7498 and Sp9509 suggests there are not large chromosomal structural variants, and that the two clones are highly similar in terms of chromosome organization. The observed differences between two genome assemblies for the same Sp7498 clone could be explained by several hypotheses.

First, genome assemblies are subject to variation depending on sequencing technology, including read lengths and error profiles. Similarly, the choice of long-read genome assembler and subsequent polishing steps can influence assembly outcome. Additionally, the method of long-range scaffolding (e.g. Hi-C, linked reads, BioNano optical mapping) can alter the scaffolding and error-correction of the final assembly. Both the Sp7498_HiC and Sp9509 assemblies were produced using Oxford Nanopore long-reads generated with MinION 9.4.1 flow cells, then assembled and polished similarly using miniasm, medaka, and RACON. Whereas the Sp9509 assembly was scaffolded with BioNano optical mapping, the Sp7498_HiC assembly was scaffolded using Phase Genomics Hi-C and their proprietary Proximo software. On the other hand, the Sp7498_PNAS assembly was generated with PacBio Sequel I reads, assembled with FALCON (Chin *et al*., 2016), and ordered using BAC-FISH. Oxford Nanopore and PacBio Sequel reads have different biases during nucleotide sequencing, which certainly could impact the full-length assembly and accurate polishing of homopolymer repeats and satellite repeats in particular. The abundance of mismatched and short indels between Sp7498_HiC and Sp7498_PNAS could be explained in part by these sequencing technology biases. Given the high degree of synteny between Sp7498_HiC and Sp9509, and mutual disagreements with large structural variations like chromosome arm inversions in Sp7498_PNAS, it is likely that BAC-FISH scaffolding introduced order and orientation errors in the Sp7498_PNAS genome.

Based on the LTR retrotransposon annotations, there is slight variation in the number of LTR retrotransposons accurately assembled between the assemblies. Identifying LTRs in the original 454 pyrosequencing plus Sanger assembly (Sp7498_v3) (Wang *et al*., 2014) shows that these short- and mid-length reads led to a collapsing of retrotransposons, especially recently amplified ones, in the final assembly (Figure 3). Both PacBio (Sp7498_PNAS) and Oxford Nanopore (Sp7498_HiC and Sp9509) assemblies appear to have corrected this issue. The Sp7498_PNAS PacBio-based assembly resulted in a slightly higher number of retrotransposon annotations, though fairly similar in number and insertion time compared to Sp9509 and Sp7498_HiC. This variation between Sp7498_PNAS and Sp7498_HiC could certainly be reflective of true biological variation due to drift, though.

A second hypothesis is related to how *S. polyrhiza* plants are clonal, and reproduce exponentially, although rarely (if ever) do they flower in nature or in culture. In the presumed absence of meiotic recombination that normally occurs during sexual reproduction, *S. polyrhiza* plants across their global range exhibit low genetic variation, and intriguingly also low spontaneous mutation rates (Xu *et al*., 2019). The estimation of Xu *et al*. of the *S. polyrhiza* mutation rate is similar to that of eubacteria and unicellular eukaryotes. Some of the mismatch, indel, and structural variation content between Sp7498_HiC and Sp7498_PNAS could be explained by these infrequent spontaneous mutations that have since accumulated between different laboratories’ stocks of Sp7498 that was originally collected in Durham, North Carolina, USA by Elias Landoldt, compared to Sp9509 collected in 2002 from a population in Lotschen, Stadtroda, Germany.

Overall, the differences that we pinpoint across these three genomes (Sp7498_HiC, Sp7498_PNAS, and Sp9509) are relatively minor depending on the usage of the genome. For instance the set of gene annotations are largely the same in number. Each of these three genomes highlight that developing high-accuracy and syntenic contigs for *S. polyrhiza* is relatively simple using either PacBio or Oxford Nanopore long-reads. The low heterozygosity and low repeat content of the genome is likely responsible for yielding such long contigs regardless of long-read sequencing technology used. However, the choice of long-range scaffolding technology is the major determinant of the quality of assembly order and orientation. In this case, both BioNano optical mapping and Hi-C scaffolding resulted in similar chromosome-scale scaffolds between Sp9509 and Sp7498_HiC, both of which differed in the ordering and orientation of several large chromosomal pieces of Sp7498_PNAS.

Global analysis of the *Spirodela* proteome revealed a high abundance of proteins involved in generating energy in *Spirodela*. In addition to comparing the proteome to other species, this model system is very well suited to analyze proteomic responses to various environmental stress conditions like heat, heavy metal stress amongst others. Intriguingly, much of the highly expressed portion of the proteome is derived from chloroplast or associated with energy production. Comparisons of expanded leaf chloroplast density, size and arrangement between *Spirodela* and *Arabidopsis* show a similar overall area of chloroplasts, but the organelles are smaller and more abundant in *Spirodela*. This is perhaps related to the two-dimensional growth pattern of duckweeds, rapid proliferation, and the need to harvest light from a single plane. Chloroplasts in a related duckweed species *Lemna trisulca* are mobile in response to heavy metals (Samardakiewicz *et al*., 2015), suggesting that chloroplast dynamics in duckweeds may favor the ability to quickly spatially reorganize in response to environmental stress. It is unknown if this is a common feature of duckweeds, but could certainly be a key factor because of the high abundance of chloroplast-derived proteins that influence rapid growth. Further, the modulation and control of RuBisCO and other plastid-associated proteins is key to crop improvement (Parry *et al*., 2003), and its relatively high expression in duckweeds serves as a foundation for exploring the genetic, developmental, physiological, and regulatory mechanisms that underlie duckweed growth.

Given the portability and speed of the Oxford Nanopore MinION platform, the computational ease of generating highly contiguous genome assemblies, and the natural abundance of duckweed species across all continents except Antarctica, there is a substantial opportunity to bring genome sequencing and proteomics of duckweed into classroom settings that culminate in valuable, publishable discoveries. The Oxford Nanopore MinION sequencer has been successfully deployed in undergraduate and graduate education across disciplines (Zaaijer *et al*., 2016; Zeng & Martin, 2017), and we expect that ongoing updates to library preparation, pore lifespan, and sequencing devices will continue to drive the proliferation of classroom-based education tools. While portable nanopore sequencing devices certainly bring cutting-edge sequencing resources to laboratory and field scientists, they perhaps more importantly democratize the ability for publishable science to be performed in the hands of students from middle school to graduate school.

## Acknowledgements

We thank the Department of Biology at Washington University in St. Louis for funding the Oxford Nanopore genome sequencing and shotgun proteome of line Sp7498. AH was supported by a post-doctoral fellowship from the NSF National Plant Genome Initiative, award #1611853. We also acknowledge imaging support from the Advanced Bioimaging Center at the Danforth Plant Science Center and usage of the Leica SPX-8 acquired through an NSF Major Research Instrumentation grant (DBI-1337680).

## Author Contributions

AH, FM, RV, BCM, and TPM conceptualized the study. AH, NB, KE, RE, EM, and KM sequenced the genome. FM, NB, KE, RE, EM, and KM generated the proteome. AH and TPM assembled the genome and performed comparative genomic analyses. FM annotated the proteome and performed the proteomic analyses. KC performed confocal microscopy and chloroplast area calculations.

## Data Availability

The Sp7498_HiC genome browser is available at CoGE (https://genomevolution.org/coge/GenomeView.pl?embed=&gid=55812). All of the assemblies described here are also found at spirodelagenome.org. The raw Oxford Nanopore reads and Hi-C Illumina reads are available at BioProject XXXXXX. The raw sequence files for the MS datasets are available in the ProteomeXchange database under the submission number PXD17093 within the PRIDE repository (http://proteomecentral.proteomexchange.org/cgi/GetDataset).

## Supplemental Information

**Supplemental Figure 1:**
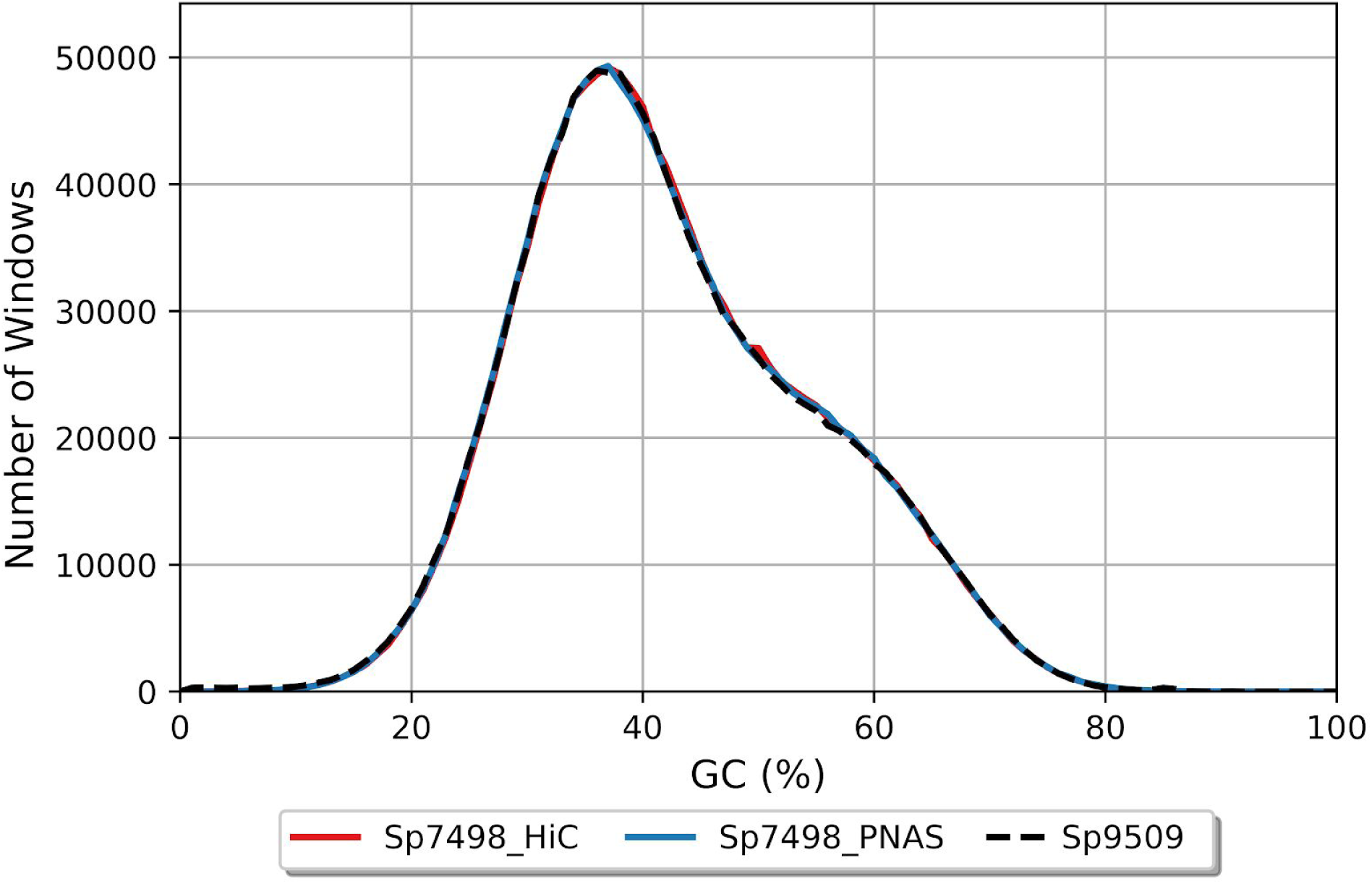
Comparison of GC content across non-overlapping 100 nt windows in the Sp7498_HiC, Sp7498_PNAS, and Sp9509 genomes.

**Supplemental Figure 2:**
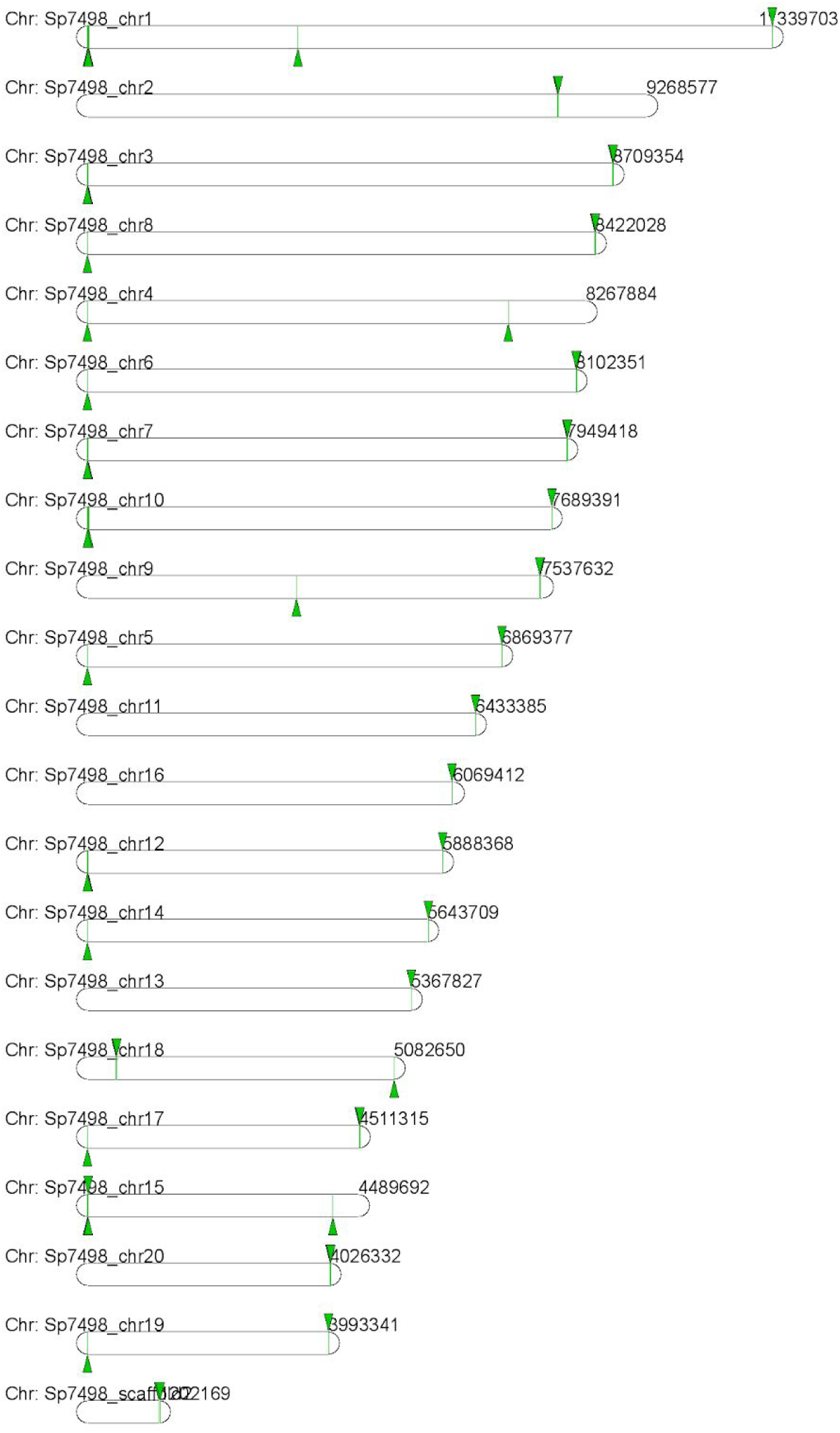
Identification and mapping of telomere repeats (5’-TTTAGGG-3’) across all 20 chromosomes of Sp7498_HiC using CoGE-BLAST. Green arrows indicate the presence of the telomere repeat.

**Supplemental Figure 3:**
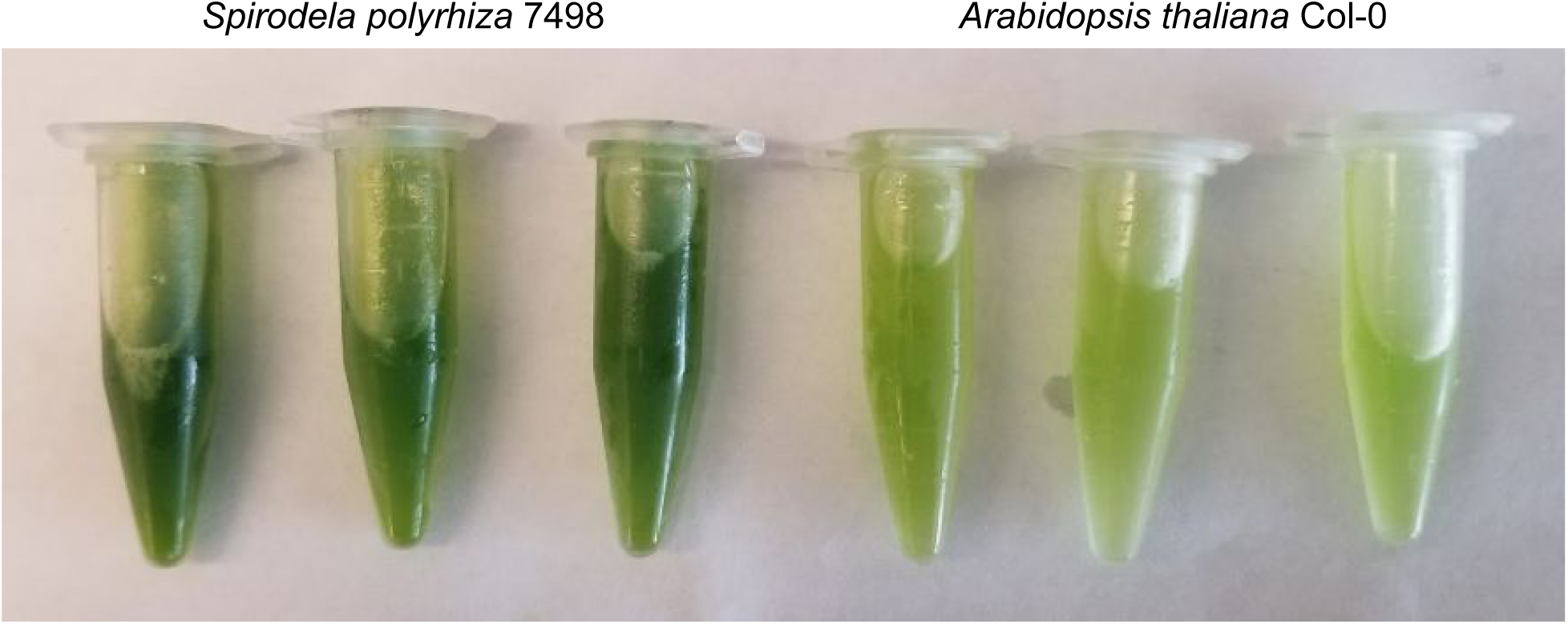
Comparison of raw protein extracts from 3 replicates of *Spirodela polyrhiza* 7498 (left) and *Arabidopsis thaliana* Col-0 (right).

